# PIDDosome-induced p53-activation for ploidy restriction facilitates hepatocarcinogenesis

**DOI:** 10.1101/2020.05.13.092486

**Authors:** Valentina Sladky, Katja Knapp, Tamas G. Szabo, Laura Bongiovanni, Hilda van den Bos, Diana C.J. Spierings, Bart Westendorp, Tatjana Stojakovic, Hubert Scharnagl, Gerald Timelthaler, Kaoru Tsuchia, Matthias Pinter, Floris Foijer, Alain de Bruin, Thomas Reiberger, Nataliya Rohr-Udilova, Andreas Villunger

## Abstract

Polyploidization frequently precedes tumorigenesis but also occurs during normal development in several tissues. Hepatocyte ploidy is controlled by the PIDDosome during development and regeneration. The PIDDosome multi-protein complex is activated by supernumerary centrosomes to induce p53 and restrict proliferation of polyploid cells, otherwise prone for chromosomal instability. PIDDosome-deficiency in the liver results in drastically increased polyploidy. To investigate PIDDosome-induced p53-activation in the pathogenesis of liver cancer, we chemically induced hepatocellular carcinoma (HCC) in mice. Strikingly, PIDDosome-deficiency reduced tumor number and burden, despite the inability to activate p53 in polyploid cells. Liver tumors arise primarily from cells with low ploidy, indicating an intrinsic pro-tumorigenic effect of PIDDosome-mediated ploidy restriction. These data suggest that hyperpolyploidization caused by PIDDosome-deficiency protects from HCC. Moreover, high tumor cell density, as a surrogate marker of low ploidy, predicts of survival of HCC patients receiving liver transplantation. Together, we show that the PIDDosome is a potential therapeutic target to manipulate hepatocyte polyploidization for HCC prevention and tumor cell density serves as a novel prognostic marker for recurrence free survival in HCC patients.

## Introduction

Hepatocellular carcinoma (HCC) represents the sixth most common cancer globally and is ranked as fourth leading cause of cancer-related mortality, and its incidence is steadily increasing (Villanueva, 2019). HCC almost exclusively develops on the background of chronic liver disease such as viral hepatitis or steatohepatitis and cirrhosis (Villanueva, 2019). Due to common late presentation with extensive disease, curative therapy is challenging and can only be provided by liver resection or transplantation (Sapisochin and Bruix, 2017). Thus, a detailed understanding of the molecular mechanisms underlying hepatocarcinogenesis is essential for disease prevention as is the identification of new therapeutic targets.

Liver cells are largely polyploid, a feature usually linked to chromosomal instability and subsequent aneuploidy, a phenomenon seen to some degree already in healthy hepatocytes (Knouse et al., 2014; Sladky et al., 2020). In most other cell types, polyploidy and chromosomal instability are considered to put cells at risk for malignant transformation (Ganem et al., 2007; Lens and Medema, 2019). Of note, more than one third of all human cancers is predicted to arise from tetraploid intermediates highlighting the significance of tight ploidy control (Zack et al., 2013).

One of the most frequently inactivated genes in HCC is the major tumor suppressor gene *TP53* which is mutated in about 30% of all patients (Lee, 2015). p53 is activated by various triggers, including DNA damage, extended mitotic timing or ploidy increases and extra centrosomes, complex structures that organize the mitotic spindle of animal cells. Active p53 restricts proliferation, induces cell death and orchestrates DNA repair to control genomic integrity and to prevent cancer (Kastenhuber and Lowe, 2017). Remarkably though, p53-induced cell death can also promote liver cancer as loss of functional parenchyma fuels compensatory proliferation in the presence of DNA damage (Qiu et al., 2011). Moreover, p53-associated activation of p21-induced cell cycle arrest can allow the survival of cells with altered genomes (Jackson et al., 2012; De La Cueva et al., 2006; Wang et al., 1997). Thus, p53 activation is not unambiguously tumor suppressive and its effects on carcinogenesis are clearly context dependent (Qiu et al., 2011).

Among all the different signals activating p53, it is the presence of supernumerary centrosomes that indicate a desired but also an aberrant increase in cellular ploidy (Andreassen et al., 2001; Godinho and Pellman, 2014). Extra centrosomes can potentially promote chromosomal instability in subsequent multipolar cell divisions, priming cells for aneuploidy (Bakhoum and Compton, 2009; Godinho and Pellman, 2014; Levine et al., 2017). Consistently, aberrant centrosome numbers are frequently found in human cancer (Chan, 2011; Godinho and Pellman, 2014). The molecular mechanism linking extra centrosomes to p53 activation involves the PIDDosome, a multi-protein complex which is formed by PIDD1 and RAIDD to activate caspase-2 (Fava et al., 2017; Tinel and Tschopp, 2004), a protease with documented tumor suppressive capacity (Ho et al., 2009; Parsons et al., 2013; Puccini et al., 2013a; Ribe et al., 2012) as well as regulatory roles in liver metabolism (Kim et al., 2009; Wilson et al., 2015).

Caspases are best-known for their roles in inflammation and cell death (Van Opdenbosch and Lamkanfi, 2019). The PIDDosome and caspase-2, however, are essential to control cellular ploidy by surveillance of centrosome numbers (Fava et al., 2017). The presence of supernumerary centrosomes indicates cell cycle defects leading to polyploidization, such as aborted mitoses or incomplete cytokinesis, and is sensed by PIDD1, residing at the mother centriole (Fava et al., 2017). Formation of the PIDDosome complex leads to caspase-2 activation, which in turn inactivates the E3-ligase MDM2 that targets p53 for proteasomal degradation. This stabilizes p53 resulting in transcriptional induction of p21 to arrest the cell cycle (Fava et al., 2017; Oliver et al., 2011). Hence, the PIDDosome prevents proliferation of cells amplifying centrosomes or following a tetraploidization step, both events associated with tumor initiation and progression (Ganem et al., 2007; Lens and Medema, 2019; LoMastro and Holland, 2019). As such, the PIDDosome should exert tumor suppressive functions that may explain observations of increased rates of tumor formation in mice lacking caspase-2, challenged by oncogenic drivers such as MYC or ERBB2, or concomitant ATM loss (Ho et al., 2009; Manzl et al., 2012; Parsons et al., 2013; Puccini et al., 2013b).

We recently showed that the PIDDosome controls physiological polyploidization in the liver (Sladky et al., 2020). Hepatocytes utilize this multi-protein complex to control the upper limit of liver ploidy during development and regeneration (Sladky et al., 2020). In the liver, expression of the centrosome-sensing component PIDD1 and the executioner component caspase-2 is coupled to proliferation, as it is controlled by antagonizing members of the E2F transcription factor family, known for their essential role in ploidy control (Chen et al., 2012; Conner et al., 2003; Pandit et al., 2012). The activating transcription factor E2F1 induces *PIDD1* and *CASP2* expression which allows induction of the PIDDosome-p53-p21 axis to restrict further proliferation of polyploid hepatocytes. This is counteracted by E2F7 and E2F8 repressing PIDDosome transcription to allow for proliferation in a polyploid state in the presence of extra centrosomes (Sladky et al., 2020). Loss of either PIDDosome component drastically increases hepatocyte ploidy. While liver function seems unaffected, the degree of aneuploidy is relatively higher as this is linked to the basal ploidy state, but not limited by PIDDosome-induced p53 function itself (Sladky et al., 2020).

Based on the above, we hypothesized that the PIDDosome pathway is not only upstream of the tumor suppressor p53 but also regulates the degree of hepatocyte polyploidy and aneuploidy in response to extra centrosomes to prevent carcinogenesis (Levine et al., 2017; Serçin et al., 2016). Hence, we investigated experimentally whether PIDDosome-mediated p53 activation limits HCC development, progression or tumor karyotype evolution. Unexpectedly, our results document a tumor promoting role for p53-dependent ploidy restriction in liver cancer and that the proliferation of tetraploid cells does not drive aneuploidy in HCC. Furthermore, our findings identify the PIDDosome as a potential drug target in HCC.

## Results

### PIDDosome-induced p53 activation facilitates DEN-induced liver tumorigenesis

The PIDDosome plays a crucial role in restricting liver ploidy during development and regeneration in hepatocytes acting upstream of p53 in response to cytokinesis failure-induced centrosome accumulation (Fava et al., 2017; Sladky et al., 2020). Thus, we wanted to test whether PIDDosome-mediated p53 activation limits tumorigenesis in the liver. To address this question, we used the DEN-driven model for chemically induced HCC in wt and mice lacking either *Casp2, Raidd/Cradd* or *Pidd1*. The livers were isolated and analyzed 10 months after DEN injection (Fig. 1A). Liver tumors, as hepatocellular adenoma and carcinoma, developed in all genotypes and in all mice (except for one) and were associated with the presence of preneoplastic lesions (foci of cellular alteration, FCA) in the surrounding non-tumorous tissue (Fig. S1A, C, data not shown). Serum levels of key hepatic parameters such as aminotransferases (ALT, AST), total bilirubin and urea were comparable between genotypes while lower cholesterol and triglycerides levels were found in PIDDosome-deficient animals (Fig. S1B), a finding in line with a proposed role of caspase-2 in *de novo* lipogenesis (Kim et al., 2018).

**Figure 1:**
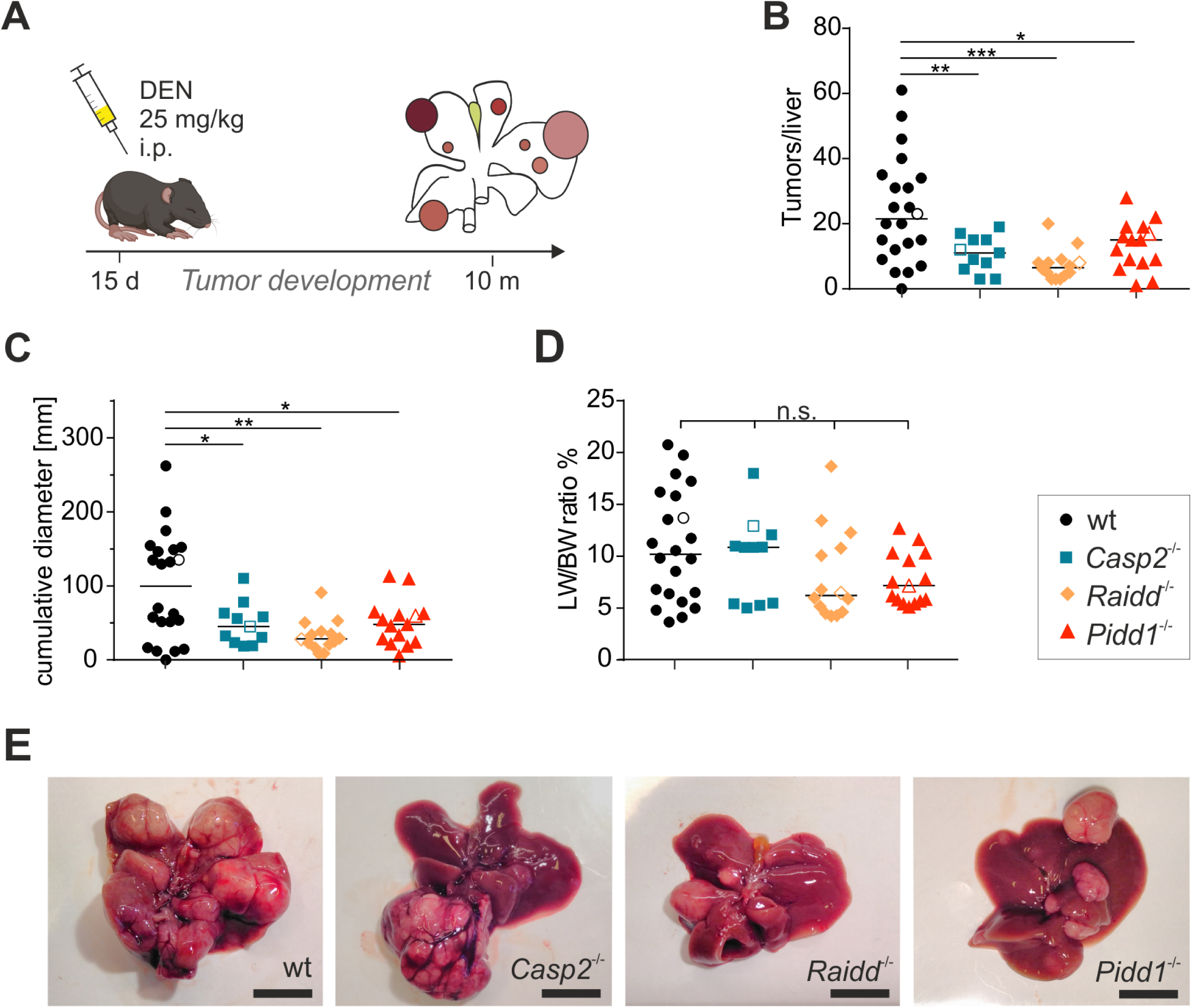
Loss of PIDDosome function reduces tumor formation in DEN-induced HCC. (A) Male wt and PIDDosome-deficient mice were injected with 25 mg/kg DEN at the age of 15 days and their livers were analyzed 10 months later. (B) Numbers of surface tumors per liver were counted. (C) The cumulative diameter of all surface tumors per liver per mouse reflects the overall tumor load. (D) Weight of the livers including tumor mass relative to the body weight (LW/BW ratio) was recorded. (E) Representative pictures of total livers with tumors of the indicated genotypes. Open symbols in graphs (B-D) represent the mice shown in (E). The line in (B-D) represents the median value of each group, statistical significance was defined as * p<0.05, ** p<0.01, *** p<0.001; scale bar in (E) represents 1 cm.

Despite similar malignancy in terms of infiltrative growth or tumor necrosis across genotypes, wt livers showed a significantly higher number of surface tumors per liver when compared to PIDDosome-deficient mice (Fig. 1B, E, data not shown). Moreover, histopathological examination of non-tumorous liver tissue revealed that the incidence of pre-neoplastic lesions (FCA) was also higher in wt mice (Fig. S1C). In line, the total tumor load was clearly higher in wt animals, reflected here by the cumulative diameter of surface tumors (Fig. 1C). Interestingly, large tumors (>8mm) tended to occur more frequently in PIDDosome-deficient mice (Fig. S1D), which likely explains the comparable liver to body weight ratios (Fig. 1D).

### DEN-induced liver tumors upregulate caspase-2 expression

Adult mouse hepatocytes express neither caspase-2 protein nor appreciable levels of *Pidd1* mRNA (Sladky et al., 2020), raising the question how the absence of the PIDDosome may affect tumorigenesis. Hence, we tested mRNA and protein levels of caspase-2 and RAIDD as well as *PIDD1* transcripts in DEN-induced HCC. Consistent with our previous results, caspase-2 protein levels were found to be barely detectable in non-tumorous, presumably healthy and largely quiescent liver tissue. Interestingly, however, caspase-2 protein was found readily expressed in all wt tumors tested (Fig. 2A). The same trends were observed for transcript levels of *Casp2* and *Pidd1* in tumor tissue (Fig. S2). Due to the lack of suitable antibodies we could not confirm increased PIDD1 protein expression in tumor tissue. Notably, we found consistent and similar RAIDD expression in both healthy liver and tumor tissue (Fig. 2A, S2).

**Figure 2:**
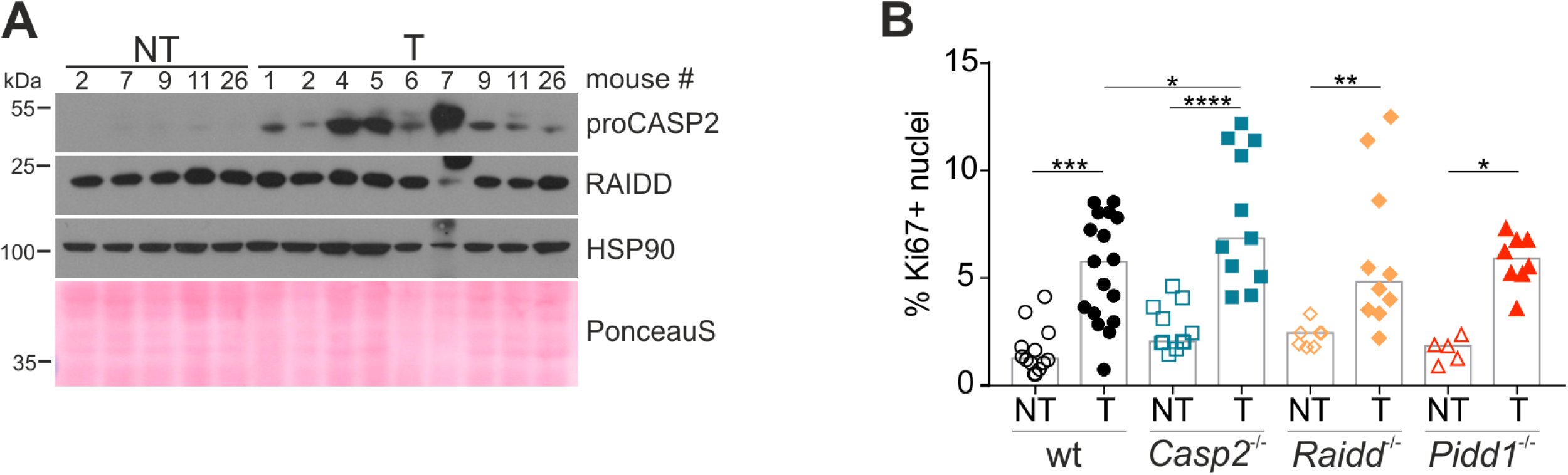
DEN-induced HCC tumors show increased proliferation and expression of caspase-2. (A) Immunoblots of murine wt non-tumorous and tumor tissue lysates were probed for CASP2 and RAIDD. HSP90 and PonceauS protein staining served as loading controls. (B) Proliferation of tumor tissue was determined by the percentage of Ki67-positive nuclei analyzed by flow cytometry. Data are represented as median, statistical significance was defined as * p<0.05, ** p<0.01, *** p<0.001, **** p<0.0001

As the PIDDosome signals upstream of p53 to limit cell cycle progression in response to extra centrosomes, we next assessed proliferation rates in liver tumors and non-tumorous liver tissue by Ki67 staining for flow cytometry and immunohistochemistry. In hepatic non-tumorous tissue, proliferation rates were low but similar in all genotypes tested. Despite appreciable differences in tumor size between wt and PIDDosome-deficient mice, Ki67 staining indicated that tumor-proliferation rates were in a comparable range, with a weak increase notable by flow cytometry only in the absence of caspase-2 (Fig. 2B). Immunohistochemistry, however, failed to confirm significant increases in Ki67^+^ tumor cells in *Casp2*^-/-^ mice (Fig. S1E). This suggests that the PIDDosome-regulated p53 response facilitates tumor initiation but most likely does not impact tumor progression. Yet, a role in limiting tumor evolution could not be excluded at that point.

### Hepatocyte ploidy defines tumor risk independent of caspase-2 and the PIDDosome

The PIDDosome functions to restrict hepatocyte ploidy and loss of either component drastically increases liver polyploidy (Sladky et al., 2020). Thus, we reasoned that the unexpected reduction in liver tumors observed in mice lacking essential PIDDosome components might be linked to its function in ploidy restriction. As DEN-induced DNA damage and compensatory proliferation on day 15, the time of DEN-injection, occurs in a mainly diploid state, the PIDDosome should not yet be activated as extra centrosomes are presumably lacking in this setting. This suggested that tumors may arise from a low ploidy state and that the ploidy increase caused by PIDDosome deficiency at the time of weaning may be protective. To test this hypothesis, we assessed the ploidy state of murine DEN-induced tumor tissue (T) and matched non-tumorous liver tissue (NT). Remarkably, in tumor tissue the numbers of binucleated and hence clearly polyploid cells per field was significantly reduced and comparable in all tumors across genotypes. In contrast, PIDDosome-deficient non-tumorous tissue tended to harbor more binucleated hepatocytes when compared to wt livers (Fig. 3A, B), consistent with our previous findings (Sladky et al., 2020). Moreover, we measured the ploidy of nuclei isolated from non-tumorous or tumor tissue which were stained with propidium iodide to assess DNA content by flow cytometry (Fig. S3A, B). Similar to binucleation tumor cell nuclear ploidy, here reflected by the percentage of octaploid nuclei, frequently found in healthy tissue (Celton-Morizur S., 2010), was clearly decreased in tumor tissue of *Casp2*^-/-^ and comparable to that found in wt mice. The same trend was observed for *Pidd1*^*-/-*^ tumors. Curiously, *Raidd*^*-/-*^ tumors did not show this effect (Fig. 3C, D). Yet, all tumors in all genotypes tested showed lower levels of polyploidy, independent of the basal ploidy state of the surrounding parenchyma (Fig. 3A). This supports the idea that HCC initiates from cells with low ploidy levels which may have a higher risk of loss of heterozygosity (LOH), and thus, are more likely to transform.

**Figure 3:**
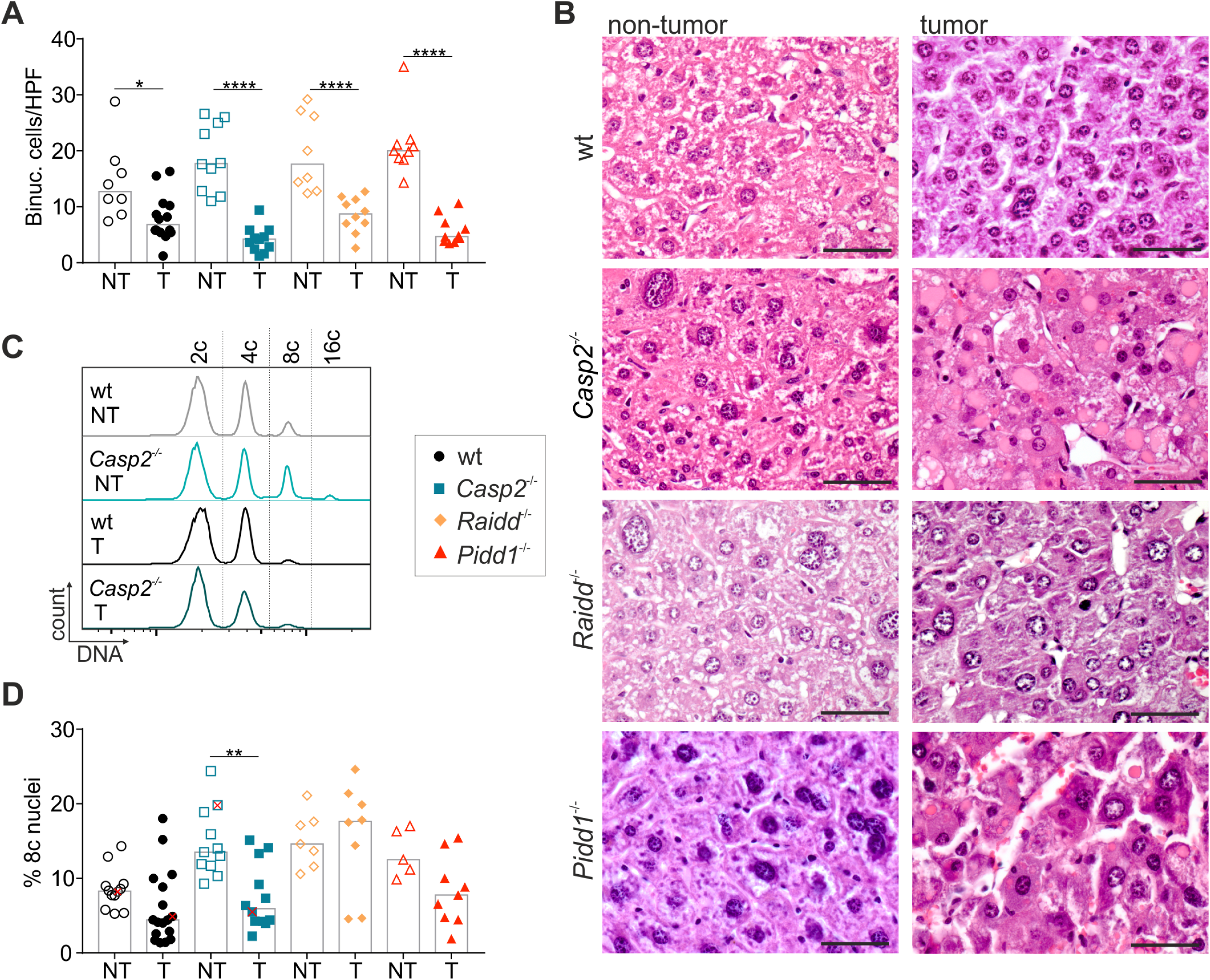
Nuclear ploidy and binucleation are reduced in HCC tumor tissue. (A) Binucleation was assessed blinded in H&E stained sections of murine non-tumor (NT) and tumor tissue (T) also shown in (B). (C) Representative histograms of nuclei stained for DNA, which were isolated from non-tumor and tumor tissue of wt and *Casp2*^-/-^ mice. These animals are marked as red “x” in the quantification shown in (D) were the degree of polyploidy is reflected as percentage of octaploid nuclei. Data are represented as median, statistical significance was defined as * p<0.05, ** p<0.01, *** p<0.001, **** p<0.0001; scale bar represents 100 µm.

### Aneuploidy is linked to basal tumor cell ploidy but not limited by Caspase-2

In the healthy liver the degree of aneuploidy increases with the polyploidy state of hepatocytes (Sladky et al., 2020). Interestingly though, caspase-2 was reported to eliminate aneuploid cancer cells *in vitro* and possibly also in patients with colorectal cancer (Dawar et al., 2016; López-García et al., 2017). Therefore, we wanted to determine the degree of copy number variation (CNV) in wt and *Casp2*^*-/-*^ liver tumors to assess the impact of PIDDosome loss on genomic stability and hence tumor evolution after transformation. To this end we subjected diploid and tetraploid nuclei isolated from wt or *Casp2*^*-/-*^ tumors to whole genome sequencing. Per tumor and ploidy state, 30 cells were pooled and subsequently sequenced to determine CNVs. Analysis of these mini bulks allows conclusions on overall aneuploidy but also intra-tumor heterogeneity as reflected by non-integer copy number states (Fig. 4A). In line with previous results in healthy liver (Sladky et al., 2020), the degree of CNV rises with the basal ploidy state but this phenomenon was independent of caspase-2 (Fig. 4A, B), and hence, presumably, the entire PIDDosome, as well as downstream p53 activity. Together, this excludes a direct role for caspase-2 in aneuploidy tolerance, at least in hepatocellular carcinoma.

**Figure 4:**
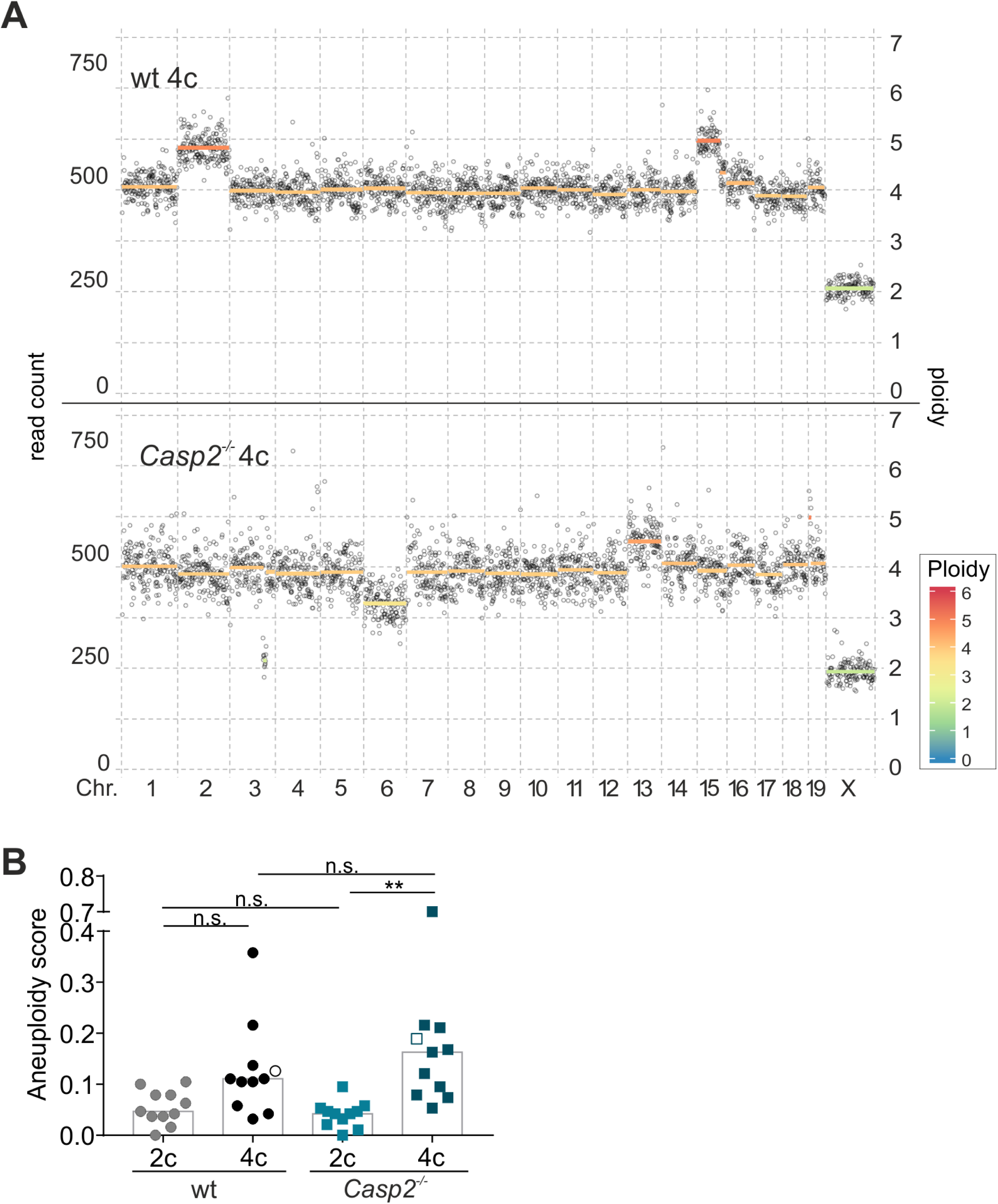
Caspase-2 has no impact on the degree of aneuploidy in HCC. Copy number variations were determined using whole genome sequencing of 30 nuclei “mini-bulks” isolated from *Casp2*^-/-^ or wt tumors. Diploid and tetraploid pools of tumor cell nuclei were analyzed separately. (A) Representative dot plots show the degree of aneuploidy and heterogeneity and are quantified in (B, empty symbols). Data are represented as median, statistical significance was defined as * p<0.05, ** p<0.01

### Caspase-2 and PIDD1 are upregulated in human HCC and correlate with poor prognosis

Since we observed increased levels of caspase-2 in murine tumor samples, we next tested the levels of each PIDDosome component in human HCC tumors and matched non-tumorous hepatic tissue. Similar to the results obtained in mice, we found caspase-2 and PIDD1 protein to be clearly elevated in human tumor tissue compared to matched non-tumorous liver. As in mice, RAIDD protein levels were comparable between tumor and healthy tissue (Fig. 5A). Based on this observation, we wondered if *CASP2* or *PIDD1* mRNA expression levels may have prognostic value in HCC. Probing the Provisional Liver Hepatocellular Carcinoma TCGA data set we found that *CASP2* and *PIDD1* transcript levels are significantly upregulated in human HCC across all disease stages. Remarkably, CASP2 mRNA levels were even higher in patients lacking functional p53. In contrast, *RAIDD* expression showed the opposite trend (Fig. 5B). In this data set, high *CASP2* expression correlated with reduced disease-free survival. This correlation, however, was not seen for *PIDD1* and *RAIDD* expression (Fig. 5C).

**Figure 5:**
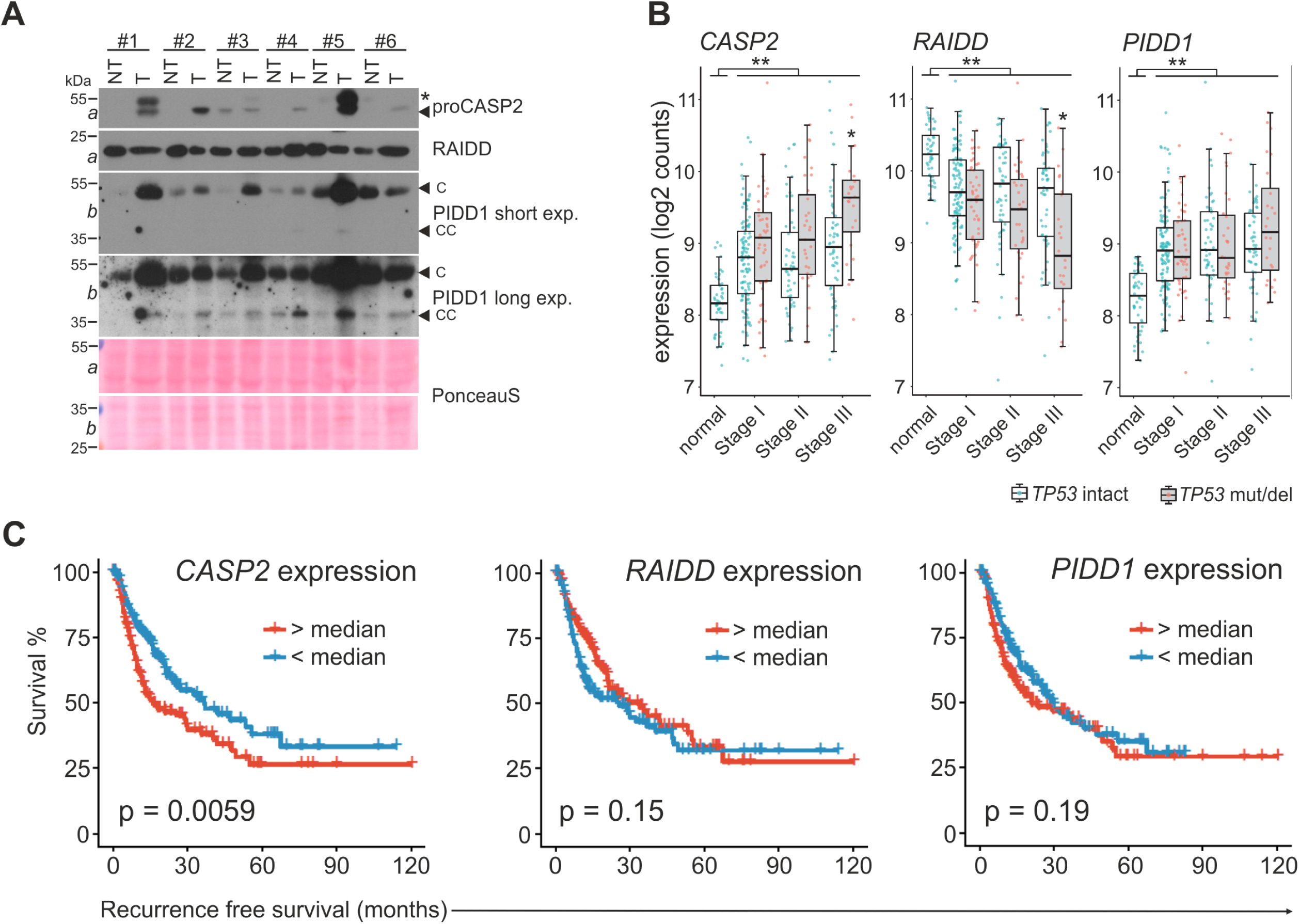
Human HCC tumors frequently upregulate CASP2 and PIDD1 expression. (A) Matched human patient non-tumor (NT) and tumor tissue (T) samples were analyzed by immunoblotting for CASP2, PIDD1 and RAIDD on membranes *a* and *b*. PonceauS protein staining serves as loading control. The asterisk refers to an unspecific band, “C” and “CC” indicate the functionally active fragments of PIDD1 generated by autoprocessing, PIDD-C and PIDD-CC. (B) mRNA expression analysis of the PIDDosome components in the TCGA Provisional LIHC data set across disease stages. Patients are divided based on the p53 mutation/deletion (mut/del) status. Data are represented as median, statistical significance was defined as * p<0.05, ** p<0.01. (C) The same data set was analyzed for recurrence-free survival based on high or low expression of CASP2, RAIDD or PIDD1 divided at the median. Statistical significance was determined using a Log-rank test.

### CASP2 and PIDD1 expression correlates with markers of proliferation

*CASP2* and *PIDD1* expression is linked to proliferation of primary hepatocytes as both are transcriptional targets of E2F transcription factors (Sladky et al., 2020). Thus, we suspected that the observed effects on expression and survival in HCC patients could be due to increased proliferation rates in these tumors which may even be increased further in the absence of p53. Consistently, expression levels of well-known proliferation markers such as Ki67 also correlated with a shorter recurrence-free survival, similar to the reduced survival noted for patients with high *CASP2*-expressing HCC (Fig. S4A). Hence, we employed bioinformatics analyses of the same TCGA data set to assess which genes are co-regulated with the PIDDosome in HCC. Strikingly, the hallmark pathways associated with genes co-expressed with *CASP2* were mostly related to proliferation, including “G2/M checkpoint”, “E2F targets” and “mitotic spindle”. Although to a minor extent, the same pattern of co-expression was found for *PIDD1*. In contrast, *RAIDD* expression was not linked to proliferation markers but, unexpectedly, was found co-regulated with genes associated with lipid metabolism (Fig. 6A, S4B). Notably, expression of *CASP2* and *PIDD1* positively correlates with other E2F target genes (Fig. 6B). Moreover, *CASP2* as well as *PIDD1* expression showed a strong positive correlation with Ki67 transcript levels while the correlation for *RAIDD* mRNA with this proliferation marker was negative (Fig. 6C-E).

**Figure 6:**
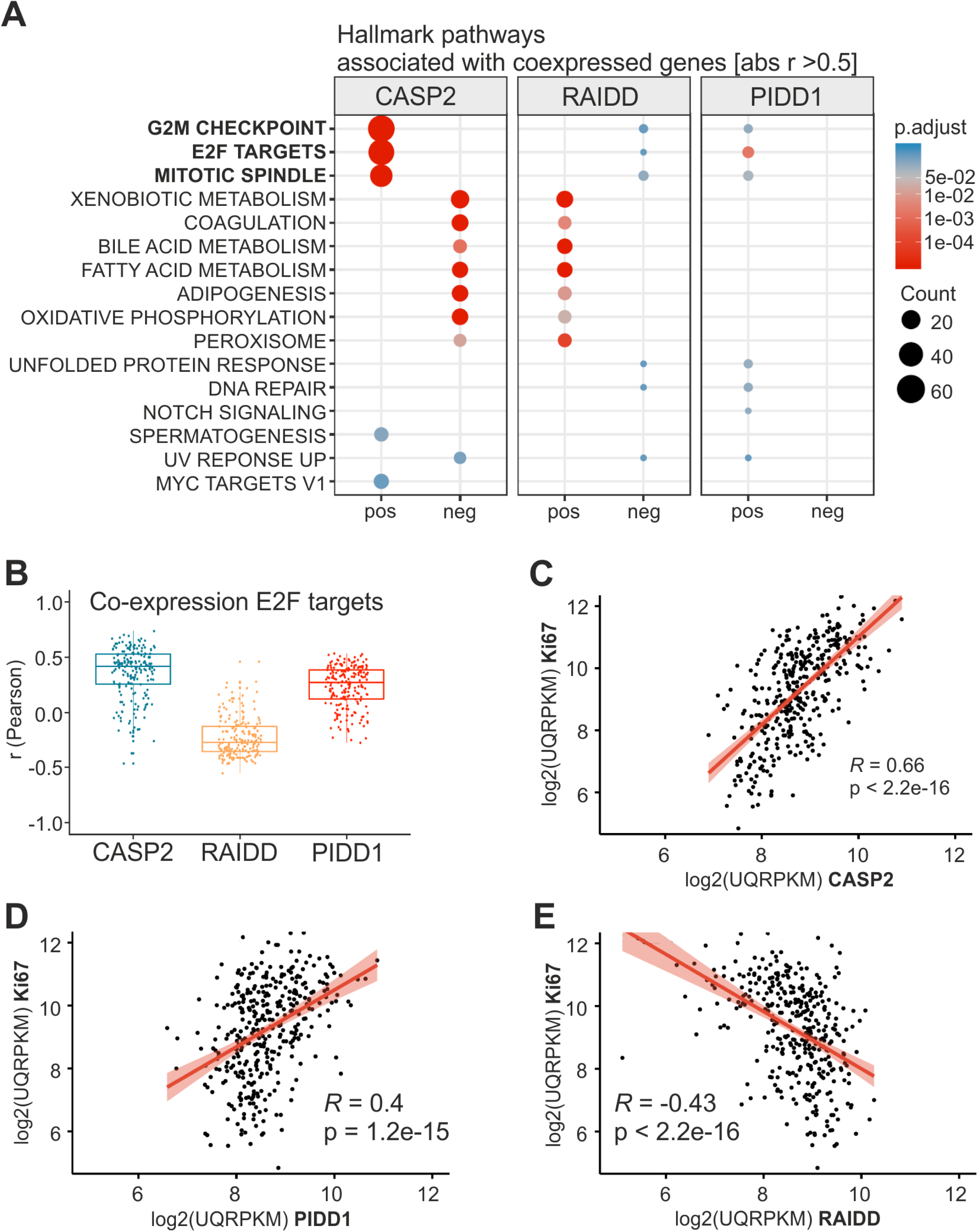
Computational analysis of the TCGA Provisional LIHC data set for expression of PIDDosome components reveals correlation with proliferation markers. (A) Genes co-expressed with the PIDDosome components were examined with respect to the associated Hallmark pathways taking positive (pos) and negative (neg) correlation into account. (B) E2F target genes expression was analyzed for correlation with expression of CASP2, RAIDD and PIDD1 using Pearson’s correlation coefficient. Data are represented as median. (C-E) Correlation (Pearson) of *CASP2, RAIDD* and *PIDD1* mRNA with transcript levels of *Ki67* in the TCGA data set.

Taken together, this suggests that the transcriptional upregulation in HCC tumors observed for *CASP2* and *PIDD1* is directly linked to the higher proliferation rates in the tumor while expression of RAIDD is uncoupled from the proliferative state. Thus, the impact of PIDDosome on hepatocarcinogenesis in mice and the recurrence-free survival in HCC patients is most likely not related to direct effects of the PIDDosome on tumor cell proliferation but rather linked to the key role of the PIDDosome controlling hepatocyte ploidy.

### Tumor ploidy differs with disease etiology and correlates with recurrence-free survival

Next we asked whether the interrelation of polyploidy and murine HCC is also seen in liver cancer patients. Therefore, we assessed the cell density of tumor and matched non-tumorous tissue of 208 patients with histologically-confirmed HCC by morphometric image analysis, as previously described (Fig. S5B-C) (Rohr-Udilova et al., 2018). Cell density, or cells per field, is an indirect, reciprocal read-out for cell size and thus ploidy. For better data visualization we used the reciprocal value of the cell density to calculate a “ploidy index” (1/cell density) for HCC tumor and non-tumorous tissue samples of each patient. Of note, the original cell density data was used to determine statistical significance and the “ploidy index” is used only for data representation. As shown before (Bou-Nader et al., 2019; Gentric et al., 2015; Toyoda et al., 2005), the etiology of the underlying liver diseases impacts on the degree of polyploidy. Non-tumorous tissue of patients with NASH or HBV infection was significantly more polyploid than HCV infected liver (Fig. 7A). Importantly, similar to our results obtained in mice and in line with previous studies (Bou-Nader et al., 2019; Gentric et al., 2015; Toyoda et al., 2005), we found that the ploidy in tumor tissue tended to be lower than in the surrounding liver. This decrease was highly significant for HCC patients with NASH. (Fig. 7A). To test whether the tumor ploidy affects disease outcome we pooled all available data on HCC tumor ploidy and divided the data according to the median HCC tumor cell density into two groups (Fig. 7B). Remarkably, the group with low HCC tumor ploidy (high cell density) showed a significantly shorter recurrence-free survival (RFS). Strikingly, multivariate analysis of age, sex, tumor size, vascular invasion, and tumor cell density clearly shows that the cell density represents an independent prognostic parameter for recurrence-free survival of HCC patients (supplemental table S2).

**Figure 7:**
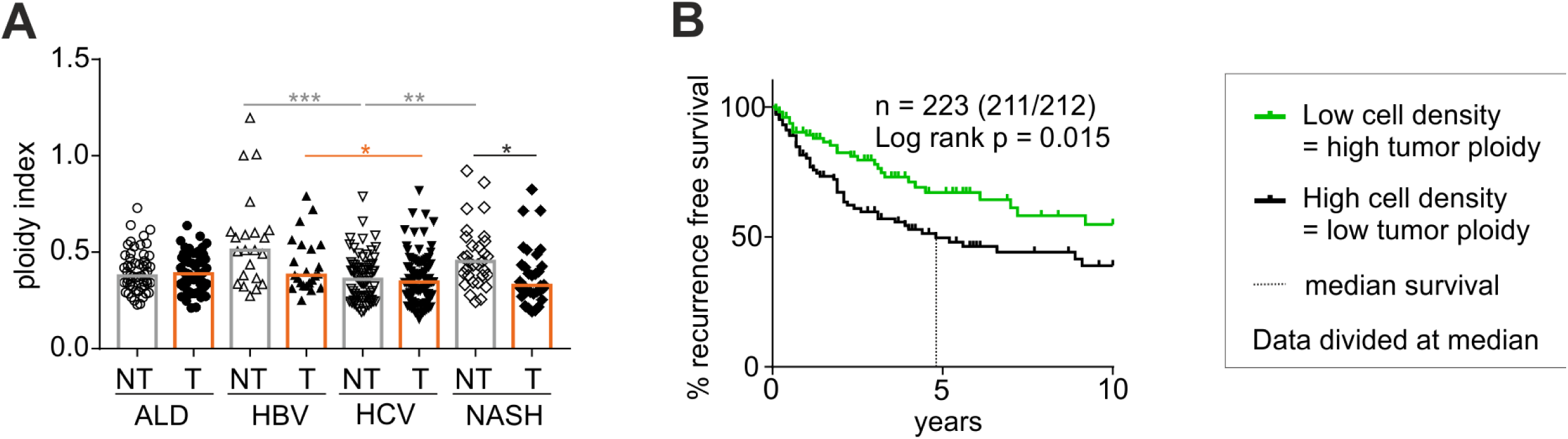
The degree of polyploidy differs between etiologies and low tumor ploidy correlates with reduced disease-free survival. (A) The ploidy index is inferred as 1/cell density of tumorous (T) and non-tumorous (NT) tissue of HCC patients with different liver diseases. ALD – alcoholic liver disease; HBV – hepatitis B virus infection; HCV – hepatitis C virus infection; NASH – non-alcoholic steatohepatitis. (B) All patients were grouped according to HCC tumor cell density as surrogate for tumor ploidy. Data were divided at the median and exact patient numbers are shown in the graphs. Median recurrence-free survival (RFS) for a period of 10 years for patients with low tumor ploidy was 4.8 years as indicated by the dashed line. For patients with high tumor ploidy the median RFS could not be defined as more than 50% of the patients survived without relapse. Patient characteristics are shown in supplemental table S1. Statistical significance was defined as * p<0.05, ** p<0.01, *** p<0.001

Together, these findings in patients with HCC support the idea that low ploid tumors are more proliferative and potentially more aggressive, increasing the risk of disease recurrence and resulting in impaired survival also after liver transplantation. As such, tumor cell density can serve as an independent prognostic marker, while *CASP2* mRNA expression serves as a surrogate of the proliferative capacity of tumor cells comparable in paucity to the well-established proliferation marker Ki67.

## Discussion

Remarkably and despite the fact that caspase-2 and the PIDDosome act upstream of the key tumor suppressor p53, we found that loss of PIDDosome-induced p53 function inhibits hepatocarcinogenesis, pointing towards an oncogenic feature of the most prominent tumor suppressor in the liver. This notion is supported by our observation that animals defective in PIDDosome function develop fewer tumors with an overall reduced tumor burden in DEN-induced HCC (Fig. 1). Notably, DEN treatment induces compensatory proliferation in response to cell death that is the central pathogenic mechanism for disease onset in this model (Qiu et al., 2011). Similar findings have been made in a mouse model of irradiation-driven lymphoma where compensatory proliferation of hematopoietic stem and progenitor cells in response to radiation-induced cell death drives malignant disease (Labi et al., 2010). So, next to the induction of cell death, p53-induced cell cycle arrest for the limitation of liver ploidy appears to be the second example where activation of this tumor suppressor can be oncogenic, albeit indirectly.

We found that *CASP2* and *PIDD1* expression is increased in human HCC as well as DEN-induced murine liver cancer (Fig. 2, 5, 6). As both genes are controlled by E2F transcription factors, their expression is clearly linked to proliferation (Sladky et al., 2020). This suggests that the effects of *CASP2* expression on patient survival are most likely secondary and non-cell autonomous as caspase-2 is transcriptionally upregulated in proliferative (tumor) tissue. As such, data correlating *CASP2* or *PIDD1* expression with patient survival need to be interpreted with caution as they most likely do not constitute markers of independent prognostic value.

Our analysis of RAIDD provides several interesting aspects. Aside from its function within the PIDDosome for liver ploidy control, it is not co-regulated with *CASP2 or PIDD1* and RAIDD-deficient mice developed the least number of tumors, albeit with the highest ploidy (Fig. 1-5). Interestingly, RAIDD expression is actually negatively linked to proliferation but is co-expressed with metabolic enzymes and this points towards a role in liver metabolism, both in the healthy liver (Sladky et al., 2020) as well as in HCC. Based on its dual adaptor domain structure, it seems to be predestined for functioning within alternative signaling complexes, which might contribute to HCC. Noteworthy here, RAIDD has originally been identified as a component in the TNF (tumor necrosis factor) receptor signaling complex (Duan and Dixit, 1997) which fuels liver carcinogenesis (Vucur et al., 2017). Future work will have to further investigate its transcriptional regulation as well as potential interactors of RAIDD to explain its role in the liver.

Several recent reports describe the impact of liver cell ploidy on hepatocarcinogenesis (Zhang, 2018 Zhang, 2018, Lin, 2020). Our findings are well in line with this recently emphasized role of high liver ploidy as a barrier against HCC, as caspase-2 or PIDDosome-deficiency significantly increases hepatocyte ploidy (Sladky et al., 2020), a phenomenon that has been overlooked before (Shalini et al., 2016). Regardless, Zhang and colleagues provide solid evidence that polyploid hepatocytes harboring tumor suppressor genes in high copy number are protected from LOH and thus, malignant transformation (Zhang et al., 2018a, 2018b). In accordance, mice harboring a liver-specific deletion of E2F8 and therefore mostly diploid hepatocytes, develop spontaneous HCC at an early age (Kent et al., 2016). This emerging hypothesis that challenges the old dogma of polyploidy fostering aneuploidy and cancer is supported by the fact that both the cellular and nuclear ploidy level of murine liver tumor tissue is reduced when compared to non-tumorous surrounding tissue (Fig. 3). Strikingly, the degree of tumor ploidy was similar between wt and the otherwise highly polyploid *Casp2*^-/-^ and *Pidd1*^-/-^ livers (Fig. 3). Equally noteworthy here, an increased occurrence of CNVs upon *Casp2* deletion or transcriptional repression, as noted in transformed mouse B cells or human colorectal carcinoma, respectively (Dawar et al., 2016; López-García et al., 2017), was neither found in the healthy liver (Sladky et al., 2020) nor in chemically induced HCC (Fig. 4). This suggests cell type specific roles of caspase-2 in the limitation of aneuploidy.

In contrast to these previous studies documenting protective effects of higher hepatocyte ploidy at the time of genotoxic challenge (Kent et al., 2016; Lin et al., 2020; Zhang et al., 2018a, 2018b), deficiency in caspase-2, PIDD1 or RAIDD leads to ploidy increases long after this DNA-damage has occurred and the DNA damage response has already ceased. At the time point of DEN injection (P15), most hepatocytes are still diploid regardless of genotype (Sladky et al., 2020; Zhang et al., 2018b). This suggests that polyploidy is not only protective at the time of mutagenesis, but can delay malignant transformation also if it occurs at a later stage, as shown in our study. This observation clearly challenges current thinking about the tumor-promoting role of polyploidization in cancer. A possible explanation may be that, for example, a tetraploid cell that had duplicated a driver mutation at the time of weaning, now also harbors four copies of each tumor suppressor gene. As such, this gene duplication event shifts the balance in favor of the rheostat of tumor suppressors even at times long after the initial driver mutation has occurred. This may explain why tumors arise from hepatocytes of low ploidy, independent of the overall liver ploidy. As this low ploid fraction of hepatocytes is substantially reduced in the absence of the PIDDosome, the population at risk for transformation is simply smaller.

Of note, most studies on the interrelation of ploidy and HCC were performed in mouse models where hepatocyte ploidy was modified by genetic manipulation of key cell cycle or cytokinesis regulators (Kent et al., 2016; Lin et al., 2020; Zhang et al., 2018a, 2018b). However, enhanced polyploidy due to deregulated cell cycle proteins is not necessarily protective as mice overexpressing *CCNE1* show strong hyperpolyploidization but develop spontaneous liver cancer (Aziz et al., 2019). This finding challenges former results in support of the “ploidy-barrier” hypothesis. Yet, as PIDDosome-deficiency increases hepatocyte ploidy without the need for direct manipulation of the cytokinesis or cell cycle machineries our study provides inaugural evidence that the ploidy-barrier hypothesis is indeed correct.

Based on the above, it would be interesting to see if the degree of ploidy in the human liver inversely correlates with HCC frequency. However, as HCC in humans is usually associated with an underlying disorder that often changes hepatocyte ploidy, such a relationship will not easily be established (Bou-Nader et al., 2019; Gentric et al., 2015; Toyoda et al., 2005). Yet, we found a clear correlation between HCC tumor cell density and recurrence-free survival with differences between the etiologies in our patient cohort (Fig. 7). Automated morphometric cell density analysis as surrogate for the degree of polyploidy clearly showed that low HCC tumor ploidy is associated with reduced survival and could serve as independent prognostic parameter. This indicates that, indeed, the interrelation between ploidy and HCC, seen in different mouse models, might also hold true for HCC patients, at least for those with NASH as the underlying disease.

A recent study suggested mononuclear polyploidy as marker for HCC aggressiveness, as in their cohort patients with highly polyploid, and often p53-mutated tumors, have an increased risk for recurrence (Bou-Nader et al., 2019). Of note, the patient cohort investigated by Bou-Nader *et al*. comprised substantially fewer patients (75 vs. 223) and also the types of underlying diseases differ between our studies. Together, these differences might explain why our result, at first sight, appear to contrast the findings by Bou-Nader and colleagues. Clearly, p53 loss increases polyploidy tolerance but most certainly increases tumor aggressiveness and recurrence rates also by other means. Unfortunately, we do not have the corresponding information regarding p53 status for our patient cohort at hand to test if these would show reduced RFS within the high ploidy group. Nonetheless, to investigate whether the protective effect of hepatocyte polyploidy found in mice really translates to humans, further studies will have to assess if healthy liver ploidy could serve as prognostic marker for the individual risk of HCC development. Morphometric analysis of routinely generated H&E stained liver tissue biopsies, for example from healthy donor livers used in transplant settings, could readily be used to collect data on hepatocyte ploidy in the healthy liver for prospective studies.

Taken together, we conclude that the PIDDosome indirectly impacts on chemically induced hepatocarcinogenesis by limiting hepatocyte polyploidization during postnatal liver development. However, in human patients the various diseases underlying HCC development make the relation between ploidy and transformation more complex than what can be shown in otherwise healthy mouse models. Regardless of this limitation, multiple lines of evidence suggest that caspase-2 is a promising therapeutic target. Here, we clearly demonstrate that caspase-2-mediated ploidy control impacts on HCC initiation. In addition, caspase-2 has been reported to accelerate NAFLD and NASH by induction of *de novo* lipogenesis, which can culminate in HCC (Anstee et al., 2019; Kim et al., 2018; Machado et al., 2015). Therefore, inhibition of caspase-2 for treatment of liver disease could prevent tumorigenesis both by elevated polyploidization as well as preventing the progression of hepatic steatosis. Finally, loss of caspase-2 also accelerates liver regeneration (Sladky et al., 2020), which can be critical for the outcome in transplant settings. Hence, the development of PIDDosome inhibitors may pave the way to treat a broad range of liver pathologies.

## Materials and Methods

### DEN-induced HCC

All animal experiments were approved by the Austrian Bundesministerium für Bildung, Wissenschaft und Forschung (Tierversuchsgesetz 2012, BGBl I Nr. 114/2012, 66.011/0108/-WF/V/3b/2015). Generation and genotyping of *Caspase-2*^*-/-*^, *Raidd*^*-/-*^and *Pidd1*^*-/-*^ mice were previously described (Berube et al., 2005; Manzl et al., 2009; O’Reilly et al., 2002). All mice used were males maintained on a C57BL/6N background under standard housing and enrichment conditions with a 12h/12h light/dark cycle. 15 days-old male C57BL/6N wild-type (n=22), *Caspase-2*^*-/-*^ (n=11), *Raidd*^*-/-*^ (n=14) and *Pidd1*^*-/-*^ (n=15) mice were intraperitoneally injected during daytime with 25 mg/kg body mass DEN (diethylnitrosamine; SigmaAldrich, St. Louis, MO, Cat# N0258, CAS# 55-18-5) diluted in PBS and sacrificed for organ harvesting 10 months later. The health status of the mice was monitored visually and by monthly weight measurement. Tumor load was assessed by counting the number of tumor nodules on the liver surface, by measuring the size of surfaces tumors using a caliper, and by weighting the livers including tumors. Tissue samples of liver tumors and non-tumorous liver tissue were collected and either fixed in 4% paraformaldehyde or snap-frozen in liquid nitrogen. Blood was collected at the time of harvest. Serum levels of aspartate aminotransferase (AST), alanine aminotransferase (ALT), total bilirubin, urea, cholesterol and triglycerides in the murine HCC models were measured enzymatically (Cobas analyser, Roche Diagnostics, Mannheim, Germany) as by manufacturer’s instruction.

### Isolation of human HCC samples and non-tumorous liver tissue

Tumors and surrounding non-tumorous tissue samples of patients with histologically-confirmed HCC undergoing orthotopic liver transplantation were collected at the Vienna General Hospital, Austria. The use of these histological sections for scientific research was approved by the ethics committee at the Medical University of Vienna (#2033/2017) and patients gave informed consent. For the morphometric analyses, HCC tumor tissue and corresponding adjacent non-tumorous liver tissue from 223 patients was evaluated. Clinical data of the patients are summarized in supplemental table S1. Clinical data analysis was performed in accordance with the guidelines of the local Ethics Committee at the Medical University of Vienna.

### Histopathological examination and immunohistochemistry

Histopathological analysis of H&E stained 4 µm sections of paraffin-embedded liver was performed by board-certified veterinary pathologists, Laura Bongiovanni and Alain de Bruin, Dutch Molecular Pathology Centre, Utrecht University, NL. Number and approximate size of nodules, tumor malignancy, inflammatory infiltration, tumor necrosis and binucleation were determined. Following deparaffinization and rehydration in xylol and ethanol the tissue sections of non-tumorous liver and tumor tissue were treated 10mM citrate buffer (pH 6.0) for antigen retrieval. Blocking was performed in peroxidase solution (1% H_2_O_2_/TBS) and sections were stained consecutively with primary (Rabbit monoclonal anti Ki67 (D3B5), Cell Signaling Technology, Danvers, MA, Cat# CS12202) and secondary (Biotin-SP-goat anti rabbit AffiniPure, Jackson ImmunoResearch Laboratories, West Grove, PA, Cat# 111-065-144) antibody diluted in 5% goat serum/TBS. Vectastain ABC Kit (Vector Laboratories, Burlingame, CA, Cat# PK-6100) and ImmPACT DAB (Vector Laboratories, Burlingame, CA, Cat# SK-4105) were used for amplification and development. The immune-stained sections were counterstained with hematoxylin, dehydrated and mounted. Images were acquired on an Olympus BX45 microscope using cell^B software (version 2.7, Olympus soft imaging solutions, Shinjuku, Japan). Analysis of slides and images was performed blinded.

### Mini bulk whole genome sequencing and data analysis

Isolation and sequencing of tumor and liver tissue nuclei was performed as described before (van den Bos et al., 2019). In brief, snap frozen tissues were cut into small pieces and incubated in nuclei isolation buffer (10 mM Tris-HCl pH8.0, 320 mM Sucrose, 5 mM CaCl2, 3 mM Mg(Ac)2, 0.1 mM EDTA, 1 mM DTT, 0.1% Triton X-100). Subsequently, nuclei were isolated from the tissue homogenate by gently passage through a 70 µm filter using a syringe plunger and pelleted by centrifugation. The nuclei pellet was re-suspended into PBS containing 2% bovine serum albumin and 10 µg/ml Hoechst 33358 (ThermoFisher Scientific, Waltham, MA, Cat# H3569) and 10 µg/ml propidium iodide (ThermoFisher Scientific, Waltham, MA, Cat# P1304MP). 30 nuclei per tumor were sorted into a single well for library preparation as described previously (van den Bos et al., 2019). Sequencing was performed using a NextSeq 500 (Illumina, San Diego, CA). The resulting sequencing data were aligned to the murine reference genome (GRCm38) using Bowtie2 (v2.2.4) (Langmead and Salzberg, 2012). To analyze copy number variation the R package AneuFinder (v1.10.1) was used (Bakker et al., 2016). Following GC-correction, blacklisting of artefact-prone regions (extreme low or high coverage in control samples) and mappability checks, libraries were analyzed using the edivisive copy number calling algorithm using 1Mb bin size, modal copy number state according to the expected ploidy state and the breakpoint detection parameter was set to 0.9. The aneuploidy score of each library was calculated as average of the absolute deviation from the expected euploid copy number per bin.

### Isolation of nuclei from snap frozen tissue

The procedure was adapted for liver and tumor tissue from Bergmann and Jovinge, 2012. Snap frozen tissue was minced with a scalpel and transferred into tissue lysis buffer (0.32 M sucrose, 10mM TrisHCl pH 8, 5 mM CaCl2, 5 mM MgAc, 2mM EDTA, 0.5 mM EGTA, 1mM DTT) and homogenized using an Ultra-Turrax probe homogenizer (IKA, Staufen, Germany) at 20,000 rpm for 12 sec. To extract nuclei from intact cells, the homogenate was passed three times through a 20 g needle and remaining tissue pieces were removed by passages through 100 µm and 70 µm cell strainers. Nuclei were sedimented by centrifugation at 700 g for 10 minutes at 4 °C and resuspended in 25 ml sucrose buffer (2.1 M sucrose, 10mM TrisHCl pH 8, 5 mM MgAc, 1mM DTT). For high density centrifugation in order to remove cell debris and enrich for nuclei, BSA-coated ultracentrifugation tubes prefilled a with 10 ml sucrose buffer cushion were carefully overlaid with the dissolved nuclei pellet and centrifuged for 60 minutes at 13,000 g (Beckman, Brea, CA, Avanti S-25, JA25.50 rotor). The supernatant was discarded and the isolated nuclei were dissolved in 1 ml nuclei buffer (0.44 M sucrose, 10 mM TrisHCl pH 7.4, 70 mM KCl, 10 mM MgCl2).

### Flow cytometry

3 × 10^6^ nuclei in nuclei isolation buffer were centrifuged at 700 g for 10 minutes fixed by adding 100 µl BD Cytofix/Cytoperm solution (BD Biosciences, Cat# 554714) while vortexing and incubated for 15 minutes on ice. After a washing step with 2 % BSA in PBS, nuclei were resuspended in 100 µl Perm/Wash buffer (BD Biosciences, Franklin Lakes, NJ, Cat# 554714, prediluted 1:10 in A.D.) and incubated for 10 minutes at room temperature. Following a washing step with Perm/Wash the nuclei were incubated with A488-conjugated antibody recognizing Ki67 (Alexa Fluor® 488 anti-mouse Ki67 Antibody (16A8), BioLegend, San Diego, CA, Cat# 652417) diluted 1:50 in Perm/Wash for 30 min at room temperature. Nuclei were washed with 2 % BSA in PBS and resuspended with 1 µg/ml propidium iodide in PBS in order to stain DNA. The stained nuclei were directly analyzed by flow cytometry on a LSR Fortessa (BD).

### Protein isolation and immunoblotting

Snap-frozen tissue of mouse or human origin was lysed in Ripa buffer (150 mM NaCl,50 mM TRIS, 1 % NP-40, 0.5 % Sodium deoxycholate% SDS, 1 tablet EDTA-free protease inhibitors). The protein content was determined using Bradford’s reagent (BioRad, Hercules, CA, 500-0006) and 40-80 µg protein was used for western blotting (AmershamTM HybondTM - ECL nitrocellulose membranes, GE Healthcare, Chicago, IL) with a wet-transfer system (BioRad, Hercules, CA). Proteins of interest were detected using antibodies recognizing caspase-2 (rat monoclonal anti-Casp2 (11B4), Enzo Life Science, Farmingdale, NY, Cat# ALX-804-356; 1:1000), RAIDD (rabbit polyclonal anti-RAIDD, Proteintech, Rosemont, IL, Cat# 10401-1-AP, 1:1000), PIDD1 (mouse monoclonal anti-PIDD1 (Anto-1), Enzo Life Science, Cat# ALX-804-837, 1:500) and HSP90 (mouse monoclonal anti Hsp90, Santa Cruz, Dallas, TX, Cat# sc-13119, 1:10000). HRP-conjugated secondary antibodies used for detection were diluted 1:5000 (HRP-mouse anti rat, Cell Signaling Technology, Danvers, MA, Cat# CS7077S; HRP-mouse anti mouse, Dako, Agilent, Santa Clara, CA, Cat# P0616; HRP-mouse anti rabbit, Dako, Cat# P0448).

### RNA isolation and quantitative real time PCR

Snap-frozen tissue was grinded in liquid nitrogen and dissolved in 500 µl TRIzol. RNA was isolated by adding chloroform to 1/5 v/v. Samples were mixed and incubated at room temperature. After centrifugation at 12,000 g for 15 minutes at 4°C, the aqueous phase was transferred into fresh tubes and the equivalent volume of isopropanol was added, mixed and incubated for 10 minutes at room temperature. Purified RNA was sedimented by centrifugation and washed with 75 % ethanol. The dried pellets were resuspended DEPC treated A.D. and DNA was digested using RQ1 RNase-Free DNase kit according to manufacturer’s instructions (Promega, Madison, WI, Cat#: M6101). Finally, RNA was precipitated again with GlycoBlueTM Coprecipitant (ThermoFisher Scientific, Waltham, MA, Cat#: AM9515). The RNA content was quantified with a NanoDrop 1000 Spectrophotometer (ThermoFisher Scientific, Waltham, MA) and reverse transcribed into cDNA with iScript™ cDNA Synthesis Kit (BioRad, Hercules, CA, Cat#: 1708891) as stated in the manufacturer’s instructions. Quantitative real-time PCR was performed on a StepOnePlus System (ThermoFisher Scientific, Inc.). The fluorescence dye based qRT-PCR using AceQ qPCR SYBR® Green Master Mix (Vazyme, Nanjing, China, Cat#: P410) was performed according to the manufacturer’s protocol (10 µl 2× AceQ qPCR SYBR® Green Master Mix, 0.4 µl Rox 1, 0.2 µl Primer Mix, 2 µl cDNA filled up with A.D. to 20 µl). The following primers were used at a final concentration of 100 nM for detection of *Casp2, Pidd1, Raidd* and *Hprt* expression: mCasp2 fw: TCTCACATGGTGTGGAAGGT, mCasp2 fw: AGGGGATTGTGTGTGGTTCT, mPidd1 fw: TGTTCTGCACAGCAACCTCC, mPidd1 rev: TGGGATATGTCTGGGGGACT, mRaidd fw: GCTTATCGGAAGAAATGGAAGCC, mRaidd rev: GGCCTGTGGTTTGAGCTTTG (Sladky et al., 2020), mHprt fw: GTCATGCCGACCCGCAGTC, mHprt rev: GTCCTTCCATAATAGTCCATGAGGAATAAAC (Schoeler et al., 2019). All qPCR reactions were performed in duplicates. PCR conditions were as follows: 95 °C for 10 min, 40 cycles of (95 °C for 15 s and 60 °C for 60 s) and a melting curve with 0.3 °C increment steps up to 95 °C for 15 s. Results for gene of interest were normalized to expression of housekeeping gene HPRT and relative expression was calculated using the ^ΔΔ^Ct method.

### Data analysis

Statistical analyses were performed by Prism 7.0.0 (GraphPad Software, San Diego, CA). Student’s t-test was used for comparison of two groups and one-way or two-way ANOVA with multiple comparison correction (Sidak-Holm) was employed if more than two groups were compared. The significance levels are stated in the respective figure legends.

Gene expression data of hepatocellular carcinoma samples collected for The Cancer Genome Atlas (TCGA) project (https://www.cancer.gov/tcga) was downloaded from the GDAC Firehose website (http://firebrowse.org; 28/01/2018 release) as upper quartile normalized RPKM (UQRPKM) values provided by RSEM. Clinical information on patients, including disease-free survival, was learned from the revised dataset of Liu et al. extracted recently from the TCGA database (Liu et al., 2018). For each patient, the first primary solid tumor tissue sample in alphabetical order of sample barcodes was taken into account. Kaplan-Meyer curves and log-rank statistics to compare prognosis in groups were analyzed using the survminer R package (https://CRAN.R-project.org/package=survminer). Information on the p53 mutation status was downloaded from cBioPortal (cbioportal.org).

Pairwise Pearson’s correlation coefficients of gene expression in hepatocellular carcinoma samples were downloaded from the cBioPortal for *Ki67, E2F1, CCNA2, PIDD1, RAIDD* (*CRADD*) and *CASP2* (cbioportal.org) (Gao et al., 2013). Genes with a Pearson’s r value greater than 0.5 or smaller than - 0.5 were considered as co-regulated with the target genes. The resulting gene lists were subjected to enrichment analysis with the *enricher* function of clusterProfiler in R (Yu et al., 2012). As an alternative approach, Gene Set Enrichment Analysis with all genes ranked by the r value was also carried out using clusterProfiler. Signature gene sets including transcription factor targets and GO terms were downloaded from the MSigDB database (http://software.broadinstitute.org/gsea/msigdb) using the msigdbr package (Subramanian et al., 2005).

### Cell density measurement

To assess cell density tissue arrays as shown in (Rohr-Udilova et al., 2018) were used. Nuclear staining was performed by hematoxylin. Slide images were digitized using a Pannoramic Midi Slide Scanner (3Dhistech, Budapest, Hungary). Morphometric characteristics of single nuclei were evaluated by tissue morphometric analysis of the digitized slide images using the Tissue Studio® software (Definiens, Munich, Germany). On average, 8710±3425 cells in tumor and matched non-tumorous tissue sections were analyzed per patient. The tissue sections were previously stained for tryptase to detect mast cells (Rohr-Udilova et al., 2018). Negatively stained cells were considered for further morphometric analysis. As shown in Fig. S5B-C, the software allows the detection of nuclei and cell borders to measure the cell area. To test the impact of nuclear circularity images of 39 HCC patients were reanalyzed considering only nuclei with a circularity-index greater than 7 out of 10 for survival analysis (Fig. S5A). As no difference was seen, all nuclei stained negatively for tryptase were used for further analysis to assess cell density (Fig. S5B, C). Recurrence-free survival was depicted by Kaplan-Meier curves and compared between patient groups by Log-rank Mantel-Cox test using Prism 7.0.0, GraphPad Software.

### Multivariate analysis

The contribution of cell density on the recurrence-free survival of patients undergoing transplantation for HCC was evaluated. Recurrence and/or death were considered as event while patients were censored at last clinical contact. Parameters associated with recurrence-free survival on univariate analysis were tested for their independent prognostic impact by cox regression and stepwise backward elimination. In total the analysis included n=172 patients. Calculations were performed using IBM SPSS Statistics 25 Software.

## Acknowledgements

We are grateful to K. Rossi, I. Gaggl, J. Heppke, C. Soratroi and A. Beierfuß for excellent technical assistance or animal care. We also thank, A. Strasser and T. Mak for sharing mouse models and reagents as well as S. Sprung, J. Haybäck, M. Bergmann, and G.F. Vogel for fruitful discussion and support with histology. This work was supported by the FWF-funded Doctoral College “Molecular Cell Biology and Oncology” (W1101) to VCS and the ERC-AdG “POLICE” to AV.

## Author contributions

VCS conducted and designed experiments, analysed data, prepared figures, wrote manuscript. KK performed experiments. TGS and BW analysed TCGA liver cancer data. LB and AdB performed histological analyses. HvdB, DCJS, FF conducted single-cell whole genome sequencing and analysed the results. TS, HS analysed serum parameters. TR analysed HCC data sets, enabled access to human tissue specimens. NRU performed morphometric analysis of human tissue sections and analysed HCC data sets. AV designed research, analysed data, wrote manuscript, conceived study.

## Conflict of interest statement

The authors declare no conflict of interest

Artwork was created using Biorender.com.

